# Ecological response of *Angelica dahurica* and its rhizosphere microorganisms to the treatment of *Alternanthera philoxeroides* extracts

**DOI:** 10.1101/2024.11.07.622522

**Authors:** Yongjie Huang, Yufeng Huang, Yuting Cai, Xinmeng Li, Jie Zhang

## Abstract

To explore the ecological adaptation mechanism of the medicinal plant *Angelica dahurica* and its rhizosphere microorganisms to the invasion of Alternanthera *philoxeroides*. This study analysed the physiological and biochemical indexes of *A. dahurica* seedlings and soil enzyme activities under a series concentration of root, stem and leaf extracts from *A. philoxeroides.* Furthermore, high-throughput sequencing technology was used to detect the rhizosphere microorganism diversity of *A. dahurica.* Results showed that with the increase of extract concentration and the extension of treatment time, the activity of antioxidant enzymes (SOD, POD, CAT) in seedlings increased and then decreased, and the inhibitory effect of root extract was the most significant. The extract effect on soil enzyme activity followed the descending sequence: root ˃ stem ˃ leaf. The activities of soil enzymes S-β-GC, S-LAP and S-ALP showed an upward trend with the increase of extract concentration, and the highest enzyme activity was found at 80 g/L. *Proteobacteria*, *Acidobacteria*, *Bacteroides* and *Sphingomonas* were the dominant species of rhizosphere soil bacteria under the treatment of each extract at phyla level. With the increasing concentration of different organ extracts, the relative abundance of *Proteobacteria*, *Bacteroides* and *Sphingomonas* gradually increased, while the abundance of *Acidobacteria* decreased. PcoA principal component analysis revealed a significant differentiation between rhizosphere bacterial community structure under the extract and control treatments. These results indicated that different concentrations of *A. philoxeroides* extract could affect the growth and soil enzyme activity of *A. dahurica* and change the community structure of rhizosphere microorganisms.

## 1. Introduction

In recent years, the rapid economic development and enhanced global connectivity have facilitated the entry of invasive alien organisms into local ecosystems through various channels, including personnel transport and logistics. As these invasive species proliferate, the risk of their spread escalates, along with the associated ecological harm. These organisms possess a remarkable ability to occupy the ecological niches of native species, thereby altering the composition and structure of invaded ecosystems (Xiao et al. 2019). *Alternanthera philoxeroides* (*A. philoxeroides*), commonly known as water peanut or water amaranth, is a perennial herb from the Amaranthaceae family, native to South America. Its strong adaptability, phenotypic plasticity, and asexual reproduction enable it to rapidly colonize diverse habitats, such as wetlands and uplands, forming monodominant communities that disrupt local ecosystems. Research has demonstrated that *A. philoxeroides* can inhibit the germination and growth of other plants by releasing allelochemicals (Zuo et al. 2012). Therefore, *A. philoxeroides* has a negative effect on crop growth and yield production in the agricultural pratices (Shen et al. 2023; Ge et al. 2018).

*Angelica dahurica* (*A. dahurica*) is a plant from the Umbelliferae family, which has dual value of medicine and food. It has been used as a food and medicine herb in China for over 1000 years. Numerous studies have reported that the root of *A. dahurica* contains volatile oils (Alkan et al. 2021), coumarin (Yang et al. 2017), amino acids (Zhang et al. 2022), and other active compounds, which are widely applied in food, health care products, spices, and skincare (Thanh et al. 2022; Vasudha et al. 2020). However, following the invasion of agricultural ecosystems by the aggressive weed *A. philoxeroides*, competition for natural resources intensifies. Moreover, it releases secondary metabolites to inhibit crop growth and development, thereby affecting crop yield and quality. Therefore, exploring the allelopathic effects of *A. philoxeroides* on the growth of *A. dahurica* is crucial for ensuring the quality and medicinal value of this herb.

Currently, research on *A. dahurica* primarily focuses on its medicinal components and pharmacological effects (Wang et al. 2016). However, studies investigating the allelopathic effects of weeds like *A. philoxeroides* on *A. dahurica* and its rhizosphere microenvironment are scarce. To elucidate the allelopathic mechanisms of *A. philoxeroides* on *A. dahurica*, we used extracts from different plant organs (roots, stems, leaves) of *A. philoxeroides* to treat *A. dahurica* seedlings, analyzing their growth and the diversity of rhizosphere soil microorganisms under these treatments. This study aims to explore (1) the effects of various organ extracts of *A. philoxeroides* on the growth of *A. dahurica* and soil enzyme activity, and (2) the ecological responses of *A. dahurica* and its rhizosphere microorganisms to the extracts of *A. philoxeroides*. The findings will provide a scientific basis for further investigating the allelopathic effects of *A. philoxeroides* on agricultural ecosystems and the field management of *A. dahurica*.

## 2. Result

### 2.1 Effect of extraction solution of *A. philoxeroides* on the growth of *A. dahurica*

In response to the extract treatments, the plant height of *A. dahurica* seedlings initially increased and then decreased with increasing extraction concentrations (Fig. 1a). The maximum increase in plant height was observed at concentrations R10, S40, and L20, with values of 4.38 cm, 6.50 cm, and 6.62 cm, respectively, significantly higher than in other treatment groups. At an extract concentration of 80 g/L, the plant height increase followed the order: S80 > CK > L80 > R80. Overall, the allelopathic effects of extracts from the roots, stems, and leaves of *A. philoxeroides* on the height changes of *A. dahurica* seedlings differed, showing the pattern: root > leaf > stem.

**Fig. 1.**
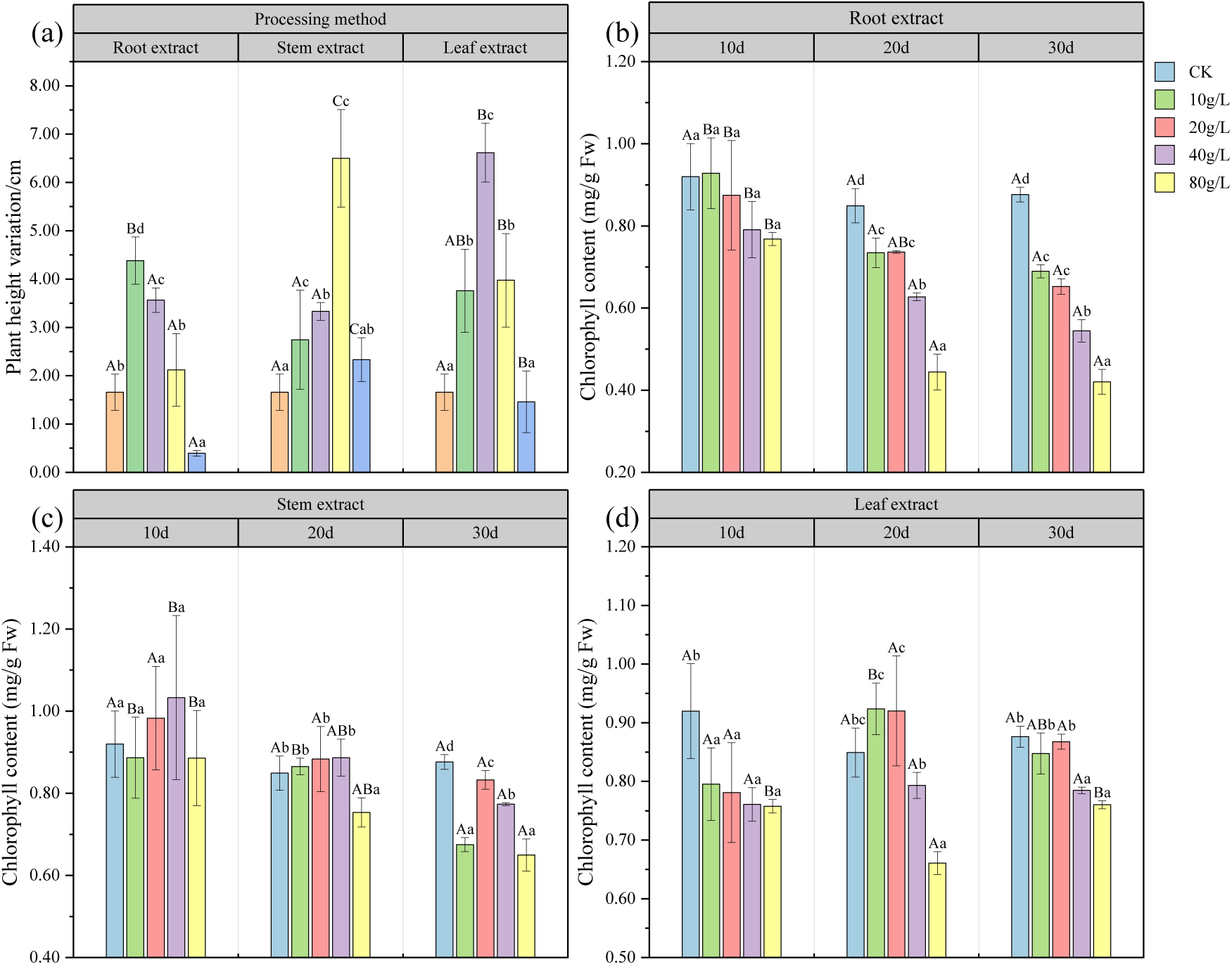
Plant height variation (**a**) and Chlorophyll content (**b, c** and **d**) of the effects of different concentrations of root, stem and leaf extract of *A. philoxeroides* on growth indexes of *A. dahurica.* Values are in bold when *p* < 0.05. Uppercase letters A, B and C indicate significant differences between the extract groups of various organs, and lowercase letter a, b and c represent the significant differences within the extract groups of different concentrations of each organ. The same below.

As treatment time extended, the chlorophyll content exhibited significant variations based on the extracted concentrations from different organs (Fig. 1b‒d). Notably, under the same concentrations of root and stem extracts, the chlorophyll content in the leaves gradually decreased over time. After 20 days of exposure to the root extract, the chlorophyll content of *A. dahurica* seedlings was significantly lower than that of the control group, with R80 showing the most pronounced reduction (Fig. 1b). On day 20, the chlorophyll content in response to stem extracts initially increased and then decreased with rising concentrations, with the 80 g/L treatment resulting in significantly lower chlorophyll levels compared to other treatment groups (Fig. 1c). Moreover, on day 30, the chlorophyll content for the S80 treatment was significantly lower than that on day 10.

On the 20th day, the chlorophyll content displayed a trend of increasing and then decreasing with increasing extract concentration, peaking at L10 (Fig. 1d). However, with prolonged treatment and rising concentrations, the chlorophyll content significantly decreased compared to the control group at concentrations exceeding 40 g/L after the 30th day. In summary, as the concentration of root extract increased and treatment duration prolonged, the chlorophyll content in *A. dahurica* seedlings generally showed a declining trend. The chlorophyll content in the leaves gradually decreased following 20 days of root extract treatment. In contrast, with stem and leaf extracts after 20 days, the chlorophyll content initially rose and then fell, with the content showing a decreasing trend after 30 days with increased extract concentrations. After 30 days of root extract treatment, chlorophyll content was significantly lower than that from stem and leaf treatments. Under leaf extract treatment, short-term treatment (≤ 10 days) reduced the chlorophyll content of *A. dahurica* seedlings, but no significant differences were observed among treatment groups. Overall, the effects of root, stem, and leaf leachate on chlorophyll content followed the order: root > stem > leaf.

### 2.2 Effect of the extract of *A. philoxeroides* on antioxidant enzyme system of *A. dahurica*

During the short-term treatment (≤ 10 days) with *A. philoxeroides* extract, the SOD activity of *A. dahurica* seedlings treated with both root and stem extracts initially decreased and then increased, peaking at 80 g/L, which was significantly higher than in other treatment groups (Fig. 2a, b). At a leaf extract concentration of 40 g/L, SOD activity was also significantly higher than the control group. However, when the concentration reached or exceeded 40 g/L, SOD activity decreased as the extract concentration increased (Fig. 2c). On day 20, the SOD activity in seedlings treated with stem extract exhibited an initial increase followed by a decrease, with significantly higher SOD activity at 10 g/L compared to other treatment groups (Fig. 2b). By day 30, as the concentrations of root, stem, and leaf extracts increased, the SOD activity in *A. dahurica* seedlings generally followed a trend of increasing and then decreasing. Both the root and stem extract treatment groups peaked at 20 g/L, while SOD activity was lowest at 80 g/L, significantly lower than in other treatment groups. The maximum SOD activity observed with leaf extract at 40 g/L was significantly higher than the control (p < 0.05). Specifically, with prolonged treatment of each extract, SOD activity gradually increased when the concentration of root extract was ≤ 20 g/L. However, at concentrations ≥ 40 g/L, SOD activity displayed a trend of increasing and then decreasing. After 30 days of treatment with high-concentration (80 g/L) stem extract, SOD activity was significantly reduced compared to the control, while there was no significant difference between leaf extract treatment and the control. Therefore, although low-concentration extracts initially promoted the growth of *A. dahurica* during short-term treatment, increasing extract concentration and treatment duration generally led to decreased SOD activity, inhibiting the growth of *A. dahurica*. In conclusion, the effects of root, stem, and leaf extracts on SOD activity in seedlings followed the order: stem > root > leaf.

**Fig. 2.**
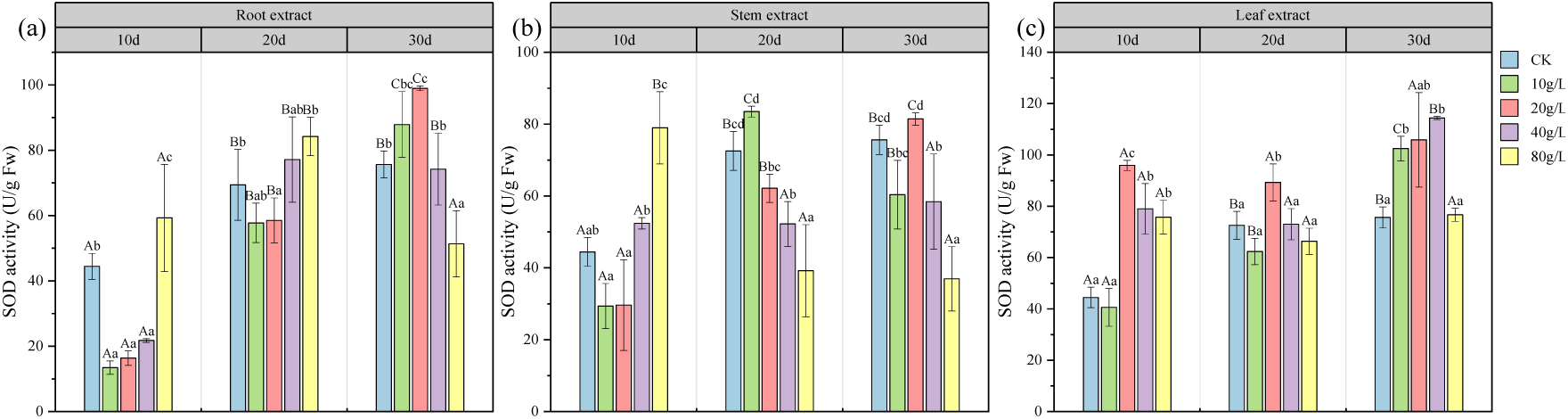
SOD activity (**a, b** and **c**) of the effects of different concentrations of root, stem and leaf extract of *A. philoxeroides* on SOD activity of *A. dahurica.* Values are in bold when *p* < 0.05

During the short-term treatment (≤ 10 d), the POD activity of *A. dahurica* seedlings increased with rising concentrations of root and leaf extracts, with POD activity at 80 g/L significantly higher than in other treatment groups (Fig. 3a, b). In contrast, the stem extract exhibited a decreasing trend followed by an increase, reaching a minimum at S20 and a maximum at S80, which was significantly higher than in other treatment groups (Fig. 3c). However, with prolonged treatment time (≥ 20 d), the overall POD activity of *A. dahurica* seedlings across all treatment groups exhibited a pattern of initial promotion followed by inhibition as extract concentrations increased. Specifically, when the concentrations of root and leaf extracts were 10 g/L, POD activity on day 30 was significantly lower than on day 10 (Fig. 3a, c). At a concentration of 80 g/L for root, stem, and leaf extracts, POD activity gradually decreased with extended treatment time and was significantly lower than after 20 days of treatment. Consequently, prolonged high-concentration treatments disrupted the antioxidant system’s balance, significantly reducing POD activity and inhibiting the growth of *A. dahurica* seedlings. The effects of root, stem, and leaf extracts on POD activity in seedling leaves were ranked as follows: root > leaf > stem.

**Fig. 3.**
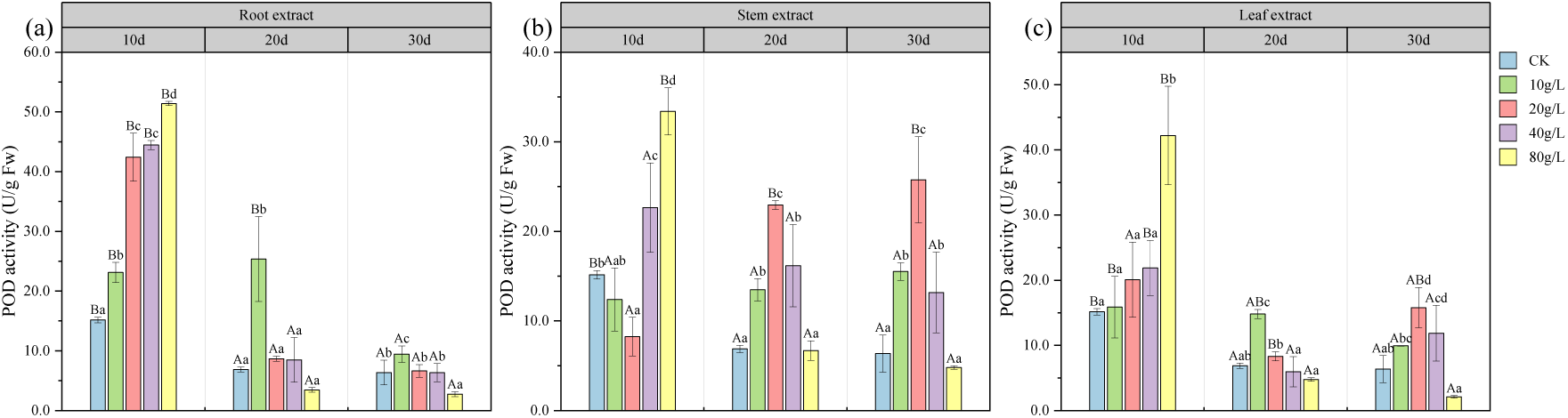
POD activity (**a, b** and **c**) of the effects of different concentrations of root, stem and leaf extract of *A. philoxeroides* on POD activity of *A. dahurica.* Values are in bold when *p* < 0.05

Overall, under the same treatment concentrations of root, stem, and leaf extracts, the CAT activity of *A. dahurica* seedlings initially increased and then decreased with extended treatment time, peaking on the 20th day (Fig. 4a, b, c). When comparing treatments at the same time point, higher treatment concentrations generally resulted in low promotion followed by high inhibition. At ≤ 10 days of treatment, CAT activity was lowest for the stem extract at 10 g/L (Fig. 4b) and for the root extract at 20 g/L (Fig. 4a), with no significant differences observed among treatment groups (Fig. 4c). After treatment with leaf extract, CAT activity gradually increased with rising treatment concentrations, with all treatment groups significantly higher than the control. On day 20, CAT activity from all extracts peaked at 20 g/L, significantly exceeding that of other groups. By day 30, for both root and leaf extracts, CAT activity first increased and then decreased with rising concentrations, reaching its lowest at 80 g/L. The influence of different organs and concentrations on *A. dahurica* seedlings exhibited a similar trend. At low concentrations (≤ 20 g/L), CAT activity gradually increased or showed a trend of initially rising and then decreasing. Conversely, for high concentrations (≥ 40 g/L), CAT activity gradually declined after 20 days of treatment, with activity at 30 days significantly lower than at 10 days of treatment. Therefore, the effects of each organ on CAT activity were ranked as follows: root > leaf > stem.

**Fig. 4.**
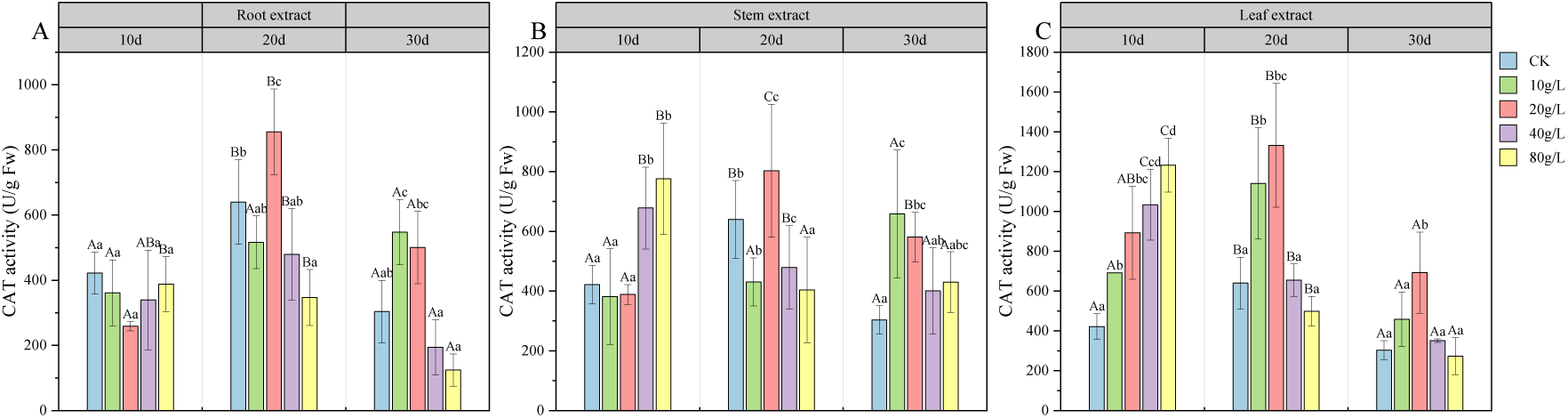
CAT activity (**a, b** and **c**) of the effects of different concentrations of root, stem and leaf extract of *A. philoxeroides* on CAT activity of *A. dahurica.* Values are in bold when *p* < 0.05.

### 2.3 Effect of *A. philoxeroides* extract on soil enzyme activities

Except for the decrease in soil S-β-GC activity observed under the high concentration of root extract (80 g/L), the activity of S-β-GC generally increased with the rising concentrations of extracts from various organs of *A. philoxeroides* (Fig. 5a). After treatment with stem and leaf extracts, S-β-GC activity increased progressively, reaching its peak at 80 g/L, which was significantly higher than the control. Under the same concentration of root, stem, and leaf extracts (≤ 40 g/L), the S-β-GC activity followed the order: root > stem > leaf. Notably, at concentrations ≤ 20 g/L, the S-β-GC activity following root extract treatment was significantly greater than that of stem and leaf treatments. Consequently, the influence of the different organ extracts on S-β-GC activity in *A. dahurica* was ranked as follows: root > stem > leaf.

**Fig. 5.**
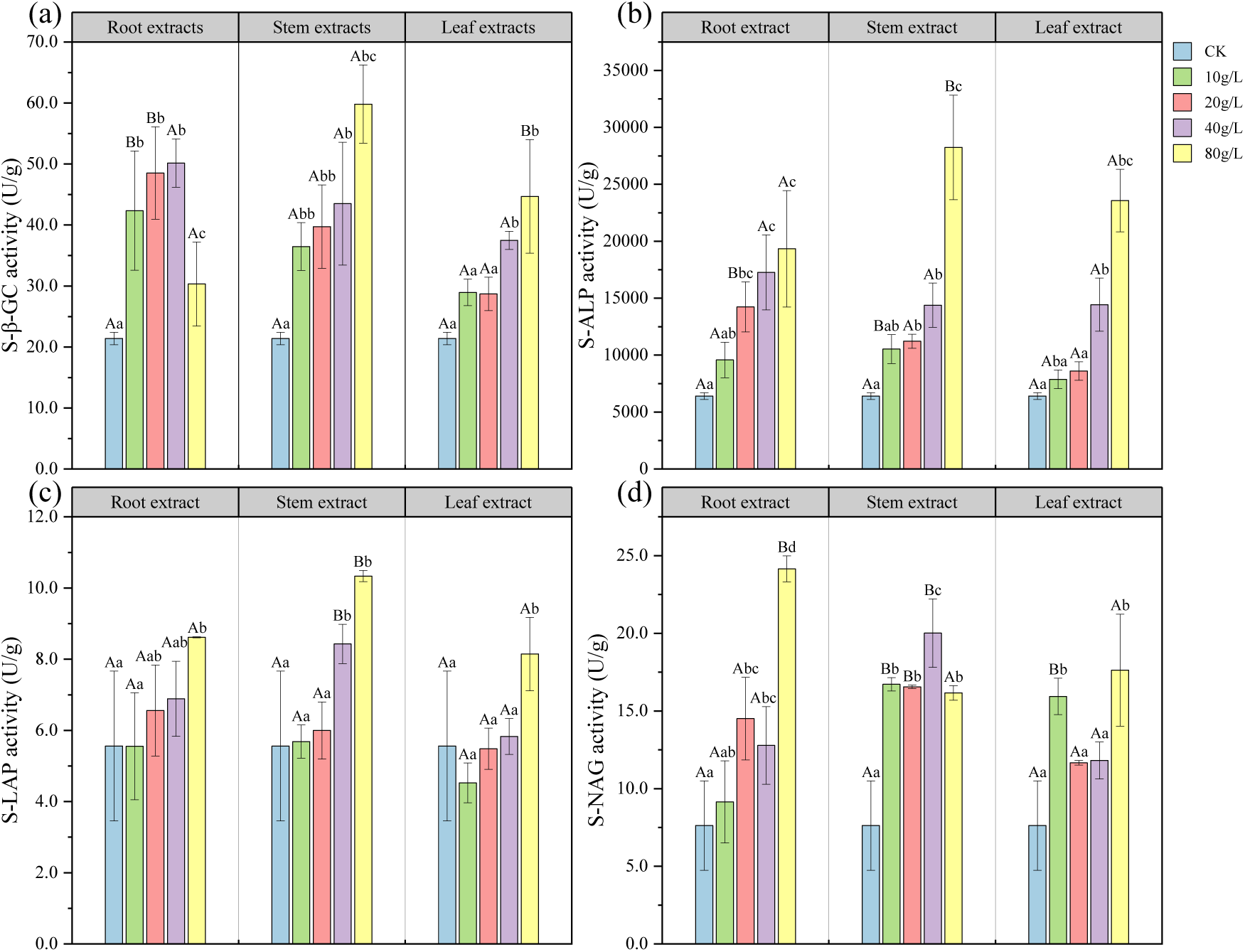
S-β-GC activity (**a**) and S-ALP activity (**b**) and S-LAP activity (**c**) and S-NAG activity (**d**) of the effects of different concentrations of root, stem and leaf extract of *A. philoxeroides* on growth indexes of *A. dahurica.* Values are in bold when *p* < 0.05

The S-ALP activity in the soil increased with higher concentrations of root, stem, and leaf extracts (Fig. 5b). When the extract concentration reached ≥ 40 g/L, S-ALP activity was significantly higher than in other treatment groups. At 80 g/L, S-ALP activity in *A. dahurica* treated with stem extract surpassed that of both root and leaf treatments, demonstrating a significant increase compared to the root extract treatment. The impact of the various organ extracts on S-ALP activity was ranked as follows: root > stem > leaf.

Overall, as the concentration of extracts from different organs of *A. philoxeroides* increased, S-ALP activity in the soil showed a clear upward trend. The highest S-ALP activity was observed at 80 g/L, significantly exceeding that of the control (Fig. 5c). The effects of different organ extracts at the same concentration on S-ALP activity varied. For instance, at concentrations ≤ 20 g/L, the differences in S-ALP activity among root, stem, and leaf treatments were not significant. However, at concentrations ≥ 40 g/L, S-ALP activity after stem extract treatment was ranked as stem > root > leaf, with stem extract treatment yielding significantly higher S-ALP activity than root and leaf treatments.

Different concentrations of extracts affected the S-NAG activity in the rhizosphere of *A. dahurica* differently (Fig. 5d). As the extract concentration increased, S-NAG activity also increased, reaching a peak at 80 g/L, significantly higher than in the control. The S-NAG activity after stem extract treatment exhibited a trend of initially increasing and then decreasing, with the highest activity recorded at a concentration of 40 g/L, significantly surpassing that of other treatments. For leaf extracts at concentrations ≥ 20 g/L, S-NAG activity increased with higher extract concentrations. Under the same extract concentration, different organs had varying effects on enzyme activity in rhizosphere soil. For example, at 10 g/L, S-NAG activity was ranked as leaf > stem > root. Conversely, at 80 g/L, the S-NAG activity under root extract treatment was significantly greater than that under stem and leaf treatments.

### 2.4 Effects of the extract of *A. philoxeroides* on the soil microorganisms in the Rhizosphere of *A. dahurica*

Analysis of the growth and soil enzyme activities of *A. dahurica* seedlings treated with various concentrations of extracts from the root, stem, and leaf of *A. philoxeroides* revealed that low concentrations (10 g/L) promoted the growth of *A. dahurica*, while high concentrations (80 g/L) inhibited normal plant growth. To further explore the ecological response of rhizosphere soil microorganisms to these two extract concentrations (10 g/L and 80 g/L), we analyzed the gene sequences of rhizosphere bacteria in the CK group and samples treated with root (G), stem (J), and leaf (Y) extracts at both concentrations. A total of 7 treatments were tested. Each treatment had 3 replicate analysis.

After sequencing 21 soil samples, we obtained a total of 1,676,641 clean reads, with each sample producing at least 79,632 reads and an average of 79,840 reads per sample. Under the influence of extracts from different organs and concentrations, the operational taxonomic unit (OTU) number in rhizosphere soil samples of *A. dahurica* initially increased rapidly with the growing number of sequences, followed by a slower rate of increase. Once the number of sequences reached a certain threshold, the dilution curves of the samples began to plateau (Fig. 6a). This curve illustrates the rate at which new species (or features) emerge with continued sampling, indicating that the sequencing depth was sufficient to accurately reflect the actual structure of the soil bacterial community.

**Fig. 6.**
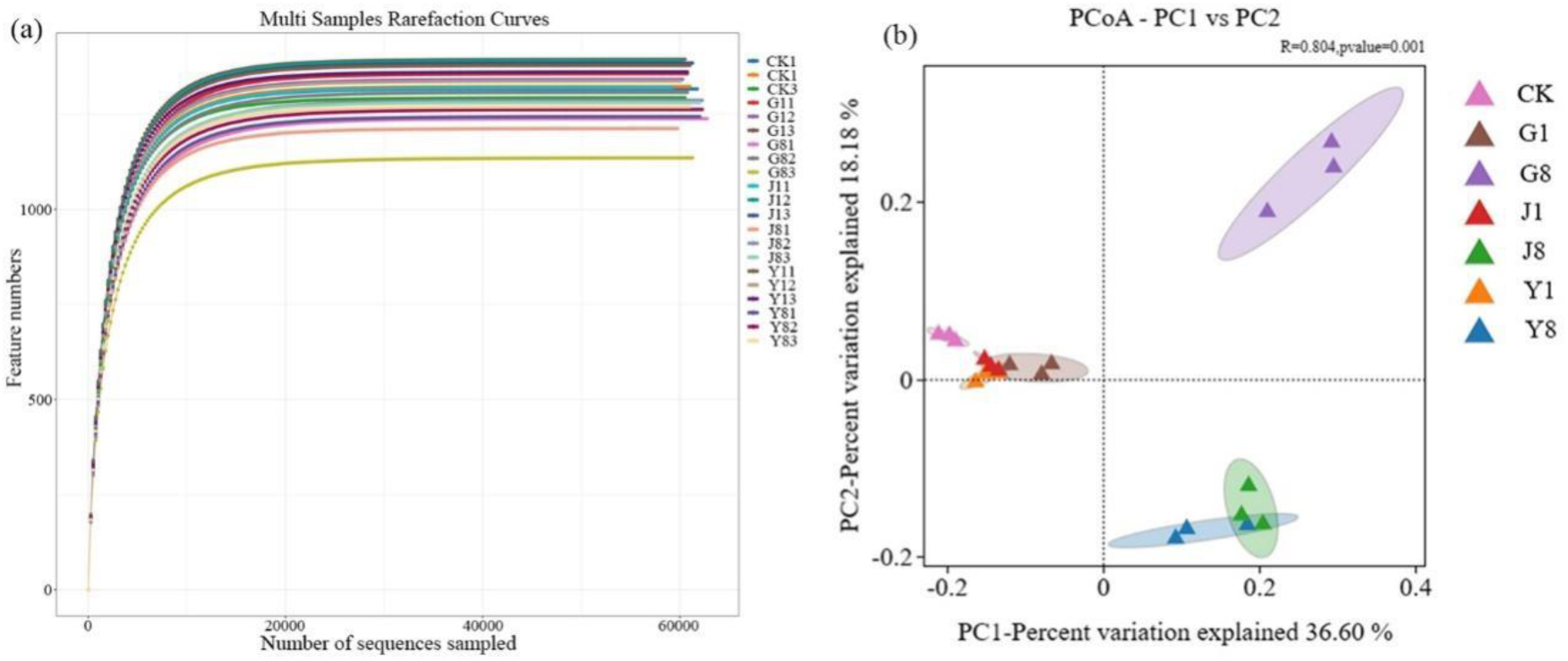
Rhizosphere soil bacterial rarefaction curves (**a**) and PCoA analysis (**b**) of the effects of different concentrations of root, stem and leaf extract of *A. philoxeroides* on rhizosphere soil bacteria of *A. dahurica*. Values are in bold when *p* < 0.05 Note: CK represents the blank control group. G1, J1 and Y1 indicated 10g/L root, stem and leaf extract treatment, respectively. G8, J8 and Y8 indicated 80g/L root, stem and leaf extract treatment, respectively. CK1‒3 stand for 3 repetitions of the set.

Alpha diversity serves as a composite index to assess both the richness and evenness of a community, thereby reflecting the species diversity within it. The Chao1 and ACE indices are indicative of the richness of fungal communities in the samples, while the Simpson and Shannon indices represent the diversity and evenness of these communities. The operational taxonomic units (OTUs) across the 21 samples ranged from 1,227 to 1,365, achieving 100% coverage (Table 1). In evaluating diversity, the results indicated that the OTUs and Shannon index values followed the order: G1 > CK > G8, J1 > CK > J8, Y1 > CK > Y8. This suggests that soil bacterial community diversity was higher under low-extract treatments. As the concentration increased, diversity among the soil bacterial communities declined. Notably, the diversity in communities treated with low-concentration extracts was significantly greater than that observed with high-concentration extracts. Additionally, the ACE estimator for low-concentration treatments was significantly higher than that for high-concentration treatments, indicating a greater abundance of soil bacterial communities in the rhizosphere soil under low-extract conditions, which decreased with increasing extract concentration.

**Table 1.** Alpha Diversity Index of Soil Bacteria in *A. dahurica* under Different Concentrations of Water Extracts from *A. philoxeroides*.

| Deal | OTUs | Shannon<br>index | Simpson<br>index | Chao1<br>index | ACE<br>index | PD_whole<br>_tree | Coverage<br>index |
| --- | --- | --- | --- | --- | --- | --- | --- |
| CK | 1310bc | 9.3116a | 0.9956b | 1310bc | 1310bc | 52.1039ab | 1.0000a |
| G1 | 1364c | 9.4854a | 0.9968b | 1364c | 1364c | 55.5203c | 1.0000a |
| G8 | 1227a | 9.2949a | 0.9967b | 1227a | 1227a | 51.5542a | 1.0000a |
| J1 | 1365c | 9.4466a | 0.9962b | 1365c | 1365c | 54.2877bc | 1.0000a |
| J8 | 1259ab | 9.0980a | 0.9930a | 1259ab | 1259ab | 53.1146ab | 1.0000a |
| Y1 | 1359c | 9.4306a | 0.9963b | 1359c | 1359c | 54.2546bc | 1.0000a |
| Y8 | 1258ab | 9.2435a | 0.9961b | 1258ab | 1258ab | 52.8399ab | 1.0000a |

Note: CK: no treatment with extract; G1: 10g/L root extract; G8: 80g/L root extract; J1: 10g/L stem extract; J8: 80g/L stem extract; Y1: 10g/L leaf extract; Y8: 80g/L leaf extract; different lower case letters in the same column: significant difference ( $p < 0.05$ ); OTUs: organism taxonomic units.

Principal Coordinate Analysis (PCoA) was employed to examine the differences in composition and structure of the soil bacterial community in the rhizosphere of *A. dahurica* seedlings treated with extracts from different organs at concentrations of 10 g/L and 80 g/L (Fig. 6b). The first principal coordinate accounted for 36.60% of the variation among the samples, effectively distinguishing the control (CK) treatment group from the rhizosphere soil bacterial community of *A. dahurica* subjected to extract treatments. This finding indicates significant differentiation in the bacterial community structure of the rhizosphere following extract application. The samples treated with low-concentration extracts (G1, J1, Y1) clustered more closely together, suggesting that the composition and structure of the bacterial communities became more similar under low-concentration treatments. In contrast, under high-concentration treatments, J8 and Y8 showed closer clustering, yet still exhibited significant differences from G8. This implies that the community composition of rhizosphere bacteria differed markedly among high-concentration treatments of root, stem, and leaf extracts (R = 0.804, p = 0.001).

### 2.5 Bacterial community composition in the rhizosphere soil of *A. dahurica*

Based on 97% similarity, the samples were classified by OTU and compared with the 16S bacterial and archaeal ribosome databases (Silva, RDP, Green Gene), revealing 28 phyla, 62 classes, 144 orders, 237 families, and 404 genera. The abundance of *Proteobacteria* and *Bacteroidota* in the soil bacterial community of *A. dahurica* showed an increasing trend with rising extract concentrations: G1 > CK > G8, CK > J1 > J8, Y1 > CK > Y8 (Fig. 7a, b). Conversely, the abundance of *Acidobacteria* decreased with increased extract concentration. At 80 g/L, the relative abundance of several phyla, including *Myxococcota*, *Methylomirabilota*, *Acidobacteriota*, *Gemmatimonadota*, *Verrucomicrobiota*, and *Crenarchaeota*, was lower compared to CK and 10 g/L treatments. Under G8 treatment, the most abundant phyla were *Proteobacteria*, *Firmicutes*, *Desulfobacterota*, and *Bdellovibrionota*, while J8 treatment exhibited aggregation of *Sumerlaeota, Abditibacteria, Patescibacteria, Cyanobacteria*, and *Deinococcota*. Notably, *Dependentiae* and *Bdellovibrionota* were the most clustered phyla under J8 treatment.

**Fig. 7.**
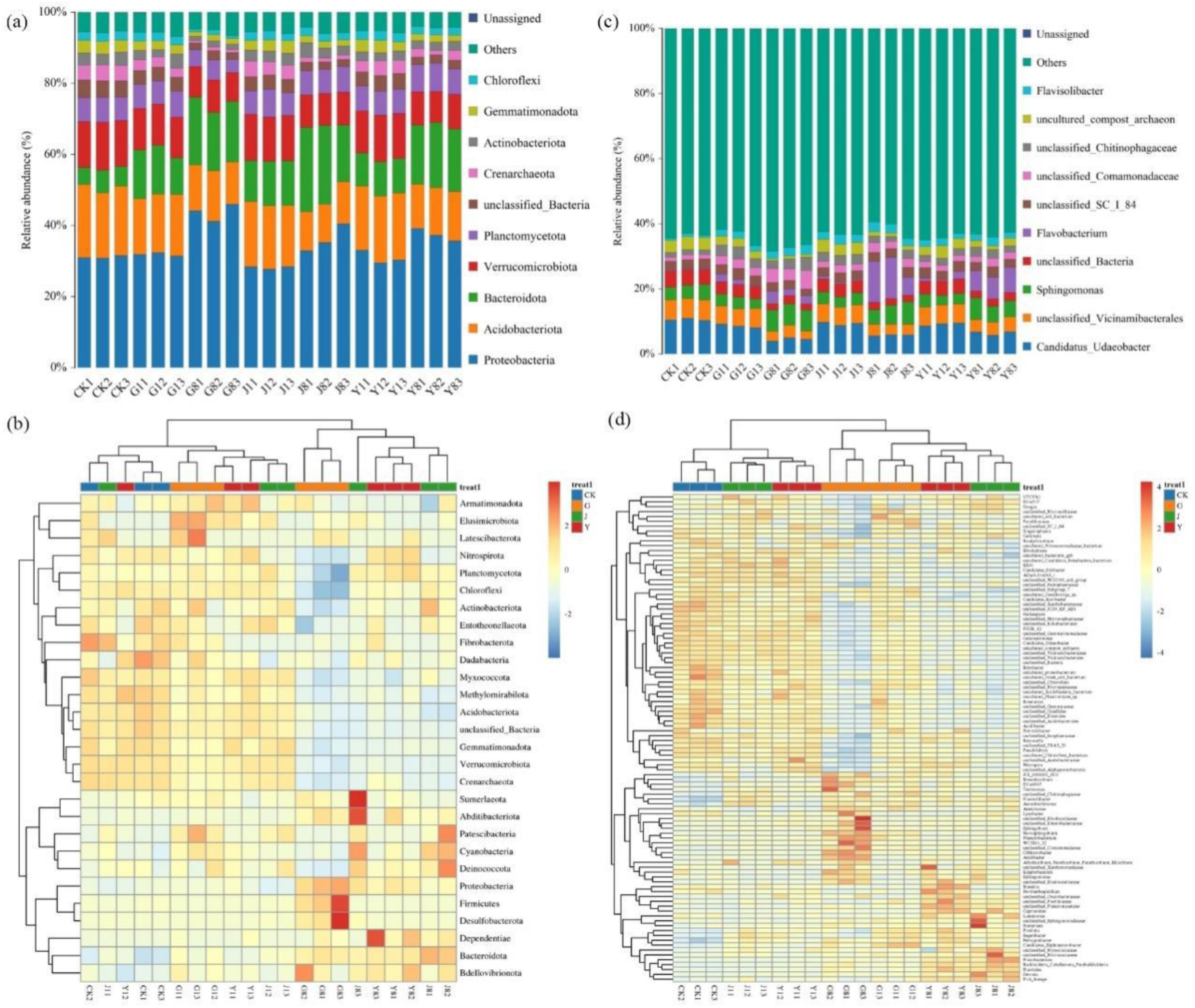
Phylum taxonomy horizontal species distribution histogram (**a**) and phylum classification level relative abundance heatmap (**b**) and genus taxonomy horizontal species distribution histogram **(c)** and taxonomic level relative abundance heatmap **(d)** of the effects of different concentrations of root, stem and leaf extract of *A. philoxeroides* on rhizosphere soil bacteria of *A. dahurica*. Values are in bold when *p* < 0.05

At the community taxonomic level, the dominant genera in the rhizosphere included *Candidatus_Udaeobacter*, *unclassified_Vicinamibacterales*, and *Sphingomonas* (Figs. 7c, d). The abundance of *Candidatus_Udaeobacter* gradually decreased with increasing extract concentration. Under G8 treatment, genera such as *Chthoniobacter*, *Novosphingobium*, *Ellin6067*, *Argillomamonas*, and *Arenimonas* showed higher aggregation. In contrast, J8 and Y8 treatments were characterized by the clustering of *Burkholderia*, *Caballeronia*, *Parburkholderia*, *Flavobacterium*, and *Sphingomonas*.

In conclusion, different concentrations of root, stem, and leaf extracts from *A. philoxeroides* significantly influenced the bacterial community structure of *A. dahurica*. Higher extract concentrations had a pronounced impact, particularly on the abundances of *Proteobacteria* and *Acidobacteria*. At the genus level, *Flavobacterium* exhibited considerable variability across different extract concentrations.

## 3. Discussion

### 3.1 The effect of *A. philoxeroides* extract on the growth and physiological indexes of *A. dahurica*

Chlorophyll plays a crucial role in plant photosynthesis, capturing light and transferring energy to the photosystems. It is vital for plant growth, regulated by stress and nutritional status. Numerous studies indicate that chlorophyll metabolism is a key indicator of plant vitality (Anne et al. 2010; Kräutler 2002). However, environmental conditions such as soil moisture, temperature, light, and stress significantly affect chlorophyll stability, leading to reduced chlorophyll content and impacting normal plant growth (Ahmadi et al. 2022; Luo et al. 2023). The findings of this study demonstrated that the reduction in chlorophyll content in *A. dahurica* seedlings was closely related to the concentration and duration of treatment with various organ extracts. Prolonged exposure to the secondary metabolites in *A. philoxeroides* extracts severely affected chlorophyll synthesis, resulting in a significant reduction in chlorophyll levels in the leaves of *A. dahurica* seedlings. Previous research has shown that allelochemicals extracted from invasive species can decrease chlorophyll and carotenoid levels, enhance the oxidative stress response in cell membranes, and induce physiological changes in treated plants, ultimately affecting the growth of native species (Andriana et al. 2018).

Reactive oxygen species (ROS) are oxygen-containing metabolites produced during plant growth. Plant aging and stressful environments can lead to irreversible oxidative damage to biomolecules, prompting cells to adopt mechanisms to prevent excessive ROS accumulation. Plants synthesize various antioxidants, including enzymatic and non-enzymatic compounds, to maintain redox homeostasis (Melicher et al. 2022; Mittler et al. 2022). Superoxide dismutase (SOD), peroxidase (POD), and catalase (CAT) are key antioxidant enzymes in the plant ROS scavenging system. The coordinated action of these enzymes effectively prevents membrane lipid peroxidation and cell damage caused by ROS, and their activity reflects the extent of plant damage (Pan et al. 2023). When treatment concentrations are too high or exposure times prolonged, levels of hydroxide ions (OH−), hydrogen peroxide (H2O2), and oxygen (O2) can increase, adversely affecting biological molecules such as amino acids, proteins, and sugars. To counter excessive free radicals and maintain physiological activity, plants regulate the activity of antioxidant enzymes, thereby balancing the antioxidant system. The results indicated that as the concentration and treatment duration of *A. philoxeroides* extracts increased, the balance of the antioxidant enzyme systems was disrupted. Over a 20-day exposure, the activities of SOD, POD, and CAT in *A. dahurica* seedlings generally followed a trend of first increasing and then decreasing. During the initial 10 days of treatment, SOD and CAT activities initially decreased and then increased with rising extract concentration, whereas POD activity consistently increased. These results suggest that shorter exposure times or lower concentrations of extracts can enhance the stress resistance of seedlings. However, prolonged exposure to high concentrations leads to an unbalanced antioxidant system and significantly reduced enzyme activity, negatively impacting the normal growth of *A. dahurica* seedlings. This finding is consistent with the experimental results of Huang et al. (2015) regarding the growth of Manila seedlings treated with water peanuts.

### 3.2 The effect of *A. philoxeroides* extract on soil enzyme activity

Soil enzymes are the most active biocatalysts in the soil ecosystem, catalyzing the decomposition of soil substances, facilitating the transformation of nutrients, and promoting nutrient cycling. Soil enzyme activity is primarily influenced by organic matter content and is often used as an indicator of microbial activity and soil fertility (Cardarelli et al. 2023). Studies indicate that soil enzyme activity responds more rapidly to changes in fertilization management, land use patterns, and planting systems than other soil indicators (Wang et al. 2023). Among these enzymes, S-β-GC hydrolyzes non-reducing sugars to release glucose, playing a crucial role in microbial metabolism and carbon (C) cycling within terrestrial ecosystems (Li et al. 2023; Fayez et al. 2019). S-NAG is another abundant enzyme, producing low molecular weight amino sugars that serve as important nitrogen sources for soil microorganisms (Cardarelli et al. 2023). S-LAP is involved in microbial nitrogen acquisition, catalyzing the hydrolysis of leucine and other hydrophobic amino acids at the N-terminus of polypeptides (Greenfield et al. 2021). S-ALP, a typical hydrolase produced by soil microorganisms and plant roots, is essential for phosphorus cycling, and its activity serves as an indicator of soil ecosystem health (Fan et al. 2020). Maričić et al. (2022) demonstrated that treating mung beans with *Urtica dioica* L. extract significantly enhanced rhizosphere soil enzyme activity of mung bean (*Phaseolus vulgaris* L.). Our study observed similar effects; the extract of *A. philoxeroides* had a marked impact on the soil enzyme activity of *A. dahurica*. Specifically, the activities of S-β-GC, S-LAP, and S-ALP all increased with higher extraction concentrations, peaking at 80 g/L. Notably, soil enzyme activity under the treatment with high-concentration (80 g/L) stem extract exceeded that of both root and leaf extracts. This enhancement may be attributed to the rich nutrient content of *A. philoxeroides*, particularly its high levels of nitrogen, phosphorus, and potassium, which improved soil fertility and subsequently increased soil enzyme activity.

### 3.3 The effect of *A. philoxeroides* extract on soil microorganisms of *A. dahurica* inter-root

Soil microorganisms perform critical processes such as oxidation, nitrification, ammoniation, and nitrogen fixation, promoting the decomposition of soil organic matter and nutrient transformation. The community structure and functional diversity of microorganisms in soil are vital factors influencing soil quality (Pan et al. 2023). Soil nutrient cycling, fertility, and productivity significantly affect the composition and structure of microbial communities (Yang et al. 2022; Partey et al. 2014). A stable microbial structure in the rhizosphere of plant roots correlates with increased species richness and diversity, enhancing plants’ stress tolerance. Studies have demonstrated a significant positive correlation between microbial diversity, community structure, microbial function, and crop yield (Shu et al. 2022). Alice et al. (2016) found that using neem (*A. zadirachta indica*) leaf water extract as a soil amendment enhanced soil microbial activity at low concentrations but inhibited it at higher concentrations. Ge et al. (2018) reported a decrease in the relative abundance of microorganisms at the genus level in treatments compared to controls. Consequently, *A. philoxeroides* may inhibit the growth of native plant species through toxic effects on soil enzyme activities and microbial communities. Our study revealed that the bacterial community composition and diversity of rhizosphere soil treated with different concentrations of root, stem, and leaf extracts of *A. philoxeroides* were significantly altered, resulting in substantial differences from the control group. The observed operational taxonomic units (OTUs), Shannon index, Chao1 index, and ACE index of soil bacteria indicated a pattern of low-concentration promotion and high-concentration inhibition. Specifically, the diversity and abundance of soil bacterial communities in the rhizosphere of *A. dahurica* were higher under low-concentration treatment of *A. philoxeroides* extract, while they declined with increased extract concentration. This difference may be attributed to the presence of nutrients in the low-concentration extraction solution that promote the growth of *A. dahurica*.

Bacteria constitute a significant portion of soil microorganisms, with *Proteobacteria* and *Acidobacteriota* being the predominant bacterial groups in terrestrial ecosystems. *Proteobacteria* are among the most abundant bacterial phyla in plant root systems, playing essential roles in ecological processes such as nitrogen cycling, organic matter decomposition, and soil remediation (Spain et al. 2009). *Bacteroidetes* are multifunctional bacteria crucial for maintaining soil functions (Kruczyńska et al. 2022), while *Acidobacteria* exhibit strong adaptability and may play unique ecological roles (Zhang et al. 2021). In our study, the dominant phyla in the rhizosphere soil of *A. dahurica* seedlings exposed to different concentrations of *A. philoxeroides* extracts were *Proteobacteria*, *Acidobacteria*, and *Bacteroidetes*. Notably, as the concentration of extracts increased, the relative abundances of *Proteobacteria* and *Bacteroidetes* rose, while the abundance of *Acidobacteria* declined. Previous studies also indicated that *Proteobacteria*, *Actinobacteria*, and *Acidobacteria* were dominant under various rhizosphere aqueous extracts of ramie (Li et al. 2023). Furthermore, spraying stevia residue extract on *Rhodopseudomonas palustris* altered soil microbial communities, significantly increasing the relative abundances of *Acidobacteria*, *Actinobacteria*, and *Proteobacteria* (Xu et al. 2016). This suggests that soil bacteria are predominantly represented by *Proteobacteria* and *Acidobacteria*, with various dominant groups coexisting due to differences in soil nutrients and other factors. At the genus level, our study found that the relative abundances of *Sphingomonas* and *Flavobacterium* in rhizosphere soil increased with higher extract concentrations. These genera possess efficient metabolic regulatory mechanisms and gene regulatory capabilities, promoting plant growth and enhancing stress tolerance (Shi et al. 2022; Feng et al. 2023). Consequently, long-term treatment with extracts from various *A. philoxeroides* organs alters the composition of soil bacterial communities, allowing bacteria with specific functions to dominate under stress. Overall, our findings indicate that *A. philoxeroides* extracts significantly impact the rhizosphere soil of *A. dahurica*, leading to substantial changes in soil bacterial community structure.

## 4. Conclusion

The high concentration of *A. philoxeroides* extract significantly inhibits the growth of *A. dahurica* seedlings, with root extract exhibiting the most pronounced inhibitory effect. This inhibition is evidenced by notable reductions in plant height, antioxidant enzyme activity, and chlorophyll content. Different concentrations of *A. philoxeroides* extract also affect the enzyme activity in rhizosphere soil and alter the microbial community structure. Specifically, soil enzyme activity is significantly enhanced under high-concentration treatment. Consequently, during the management of *A. dahurica* cultivation, it is advisable to moderate the eradication of *A. philoxeroides* to ensure the normal growth of *A. dahurica*. However, the impacts of varying concentrations of *A. philoxeroides* extracts on the chemosensory substances, medicinal components, and overall medicinal value of *A. dahurica* remain unclear, necessitating further research.

## 5. Materials and Methods

### 5.1 Preparation of Extracts

The whole plants of *A. philoxeroides* were collected, and the yellowed, withered leaves were removed. Fresh stems, leaves, and roots were washed and naturally air-dried. The dried plant parts were crushed, passed through a 60-mesh sieve to obtain dry powder, and then mixed with distilled water at varying solid-to-water ratios (10 g/L, 20 g/L, 40 g/L, and 80 g/L). The mixtures were soaked at room temperature (18‒20 °C) for 48 hours and then centrifuged at 4000 rpm for 10 minutes to obtain the supernatants containing the extracts of different concentrations. The extracted solutions were stored at 4 °C for subsequent use.

### 5.2 Experimental Design

*A. dahurica* seeds were sown in plastic pots filled with 6.0 kg of field soil (dimensions: 30 cm long, 45 cm wide, and 30 cm high). After two months of pre-cultivation, seedlings with uniform plant height and similar growth status were selected for the pot experiment. Each plastic pot (15 cm in diameter, 30 cm in height) was filled with 2.0 kg of garden soil, and three *A. dahurica* seedlings were transplanted per pot. Thirteen treatments were set up with three replicates each. After seven days of stabilization, the seedlings were treated with 100 mL of root, stem, or leaf extracts (R, S, L) at different concentrations (0 g/L, 10 g/L, 20 g/L, 40 g/L, and 80 g/L). Distilled water was used for the control group. During regular management, 100 mL of extract was applied every three days. Physiological indicators (superoxide dismutase, peroxidase, catalase, and chlorophyll content) of *A. dahurica* were measured on days 10, 20, and 30 after treatment. On the 30th day, the increase in plant height was measured, and soil samples were collected for the analysis of enzyme activity and microbial diversity in the rhizosphere using biochemical and molecular techniques.

### 5.3 Sample Collection

Leaves from the 2nd to 3rd position of *A. dahurica* were collected for the measurement of various growth indicators. Soil samples from each replicate were thoroughly mixed and treated as a single sample. After air-drying, the soil was ground, homogenized, and passed through a 2-mm sieve for enzyme activity analysis. Rhizosphere soil (approximately 1 mm thick, tightly adhered to the root system) was collected following the method described by Luo et al. (2017), without damaging plant tissues, and stored at −80 °C for DNA extraction and microbial analysis.

### 5.4 Sample Analysis

The vertical height from the root collar to the plant apex was measured with a ruler (cm). Superoxide dismutase (SOD), peroxidase (POD), and catalase (CAT) activities were measured using the NBT photoreduction method, guaiacol method, and UV absorption method, respectively (Shi, 2016). Chlorophyll content was extracted with 95% ethanol and determined by ultraviolet spectrophotometry (Wang et al., 2015). Soil β-glucosidase (S-β-GC), N-acetyl-β-D-glucosaminidase (S-NAG), leucine aminopeptidase (S-LAP), and alkaline phosphatase (S-ALP) activities were measured according to the kit instructions (Beijing Solarbio Technology Co., Ltd.).

Soil microbial samples were analyzed using 16S rRNA amplicon sequencing. DNA was extracted from soil samples and stored at −20 °C for PCR amplification using the primers F: GTGYCAGCMGCCGCGGTAA and R: CCGYCAATTYMTTRAGTTT, targeting the V4–V5 region of the 16S rRNA gene. The PCR products were purified from 2% agarose gels and sequenced using the Illumina NovaSeq 6000 platform. Primer sequences were removed, and the paired-end reads were merged to obtain the final valid data. DNA extraction, OTU clustering, species taxonomy, and sequencing analysis were outsourced to Biomarker Biotechnology Co., Ltd. (Beijing).

### 5.5 Data Processing and Statistical Analysis

Data, including physiological, biochemical parameters, rhizosphere soil enzyme activity, and microbial sequencing results, were analyzed using Excel and SPSS 19 software. One-way ANOVA was performed to compare the differences between treatments, with significance set at p < 0.05. Principal coordinate analysis (PcoA) of OTUs was performed using the R package “psych,” and all figures were generated using Origin 2023.

## Acknowledgments

This work was s supported by the Natural Science Foundation of Colleges and Universities of Anhui Province (KJ2019A0507), Wuhu Science and Technology Bureau Project (2023JC18).

## CRediT authorship contribution statement

Yongjie Huang: review, editing, and funding acquisition. Yufeng Huang: conceptualization, material preparation, data collection, experiments and writing. Yuting Cai: material preparation and experiments. Xinmeng Li: material preparation and experiments. Zhang Jie: review, editing.

## Data availability

The data underlying this article are available in Harvard Dataverse, at https://doi.org/10.7910/DVN/AVOTBT.

